# Genetic interaction between profilin and myosin II reveals a potential role for myosin II in actin filament disassembly *in vivo*

**DOI:** 10.1101/2020.04.03.023705

**Authors:** Paola Zambon, Saravanan Palani, Shekhar Sanjay Jadhav, Pananghat Gayathri, Mohan K. Balasubramanian

## Abstract

The actin cytoskeleton plays a variety of roles in eukaryotic cell physiology, ranging from cell polarity and migration to cytokinesis. Key to the function of the actin cytoskeleton is the mechanisms that control its assembly, stability, and turnover. Through genetic analyses in fission yeast, we found that, *myo2-S1* (*myo2*-G515D), a myosin II mutant allele was capable of rescuing lethality caused by compromise of mechanisms involved in actin cable / ring assembly and stability. The mutation in *myo2*-S1 affects the activation loop of Myosin II, which is involved in physical interaction with subdomain 1 of actin and in stimulating the ATPase activity of Myosin. Consistently, actomyosin rings in *myo2*-S1 cell ghosts were severely compromised in contraction upon ATP addition, suggesting that Myo2-S1p was defective in actin binding and / or motor activity. These studies strongly suggest a role for Myo2p in actin cytoskeletal disassembly and turnover, and that compromise of this activity leads to genetic suppression of mutants defective in actin cable assembly / stability.

## Introduction

Actin is a highly conserved cytoskeletal polymer forming protein that is key to a vast array of physiological processes in the three domains of life. In eukaryotes, the actin cytoskeleton plays essential roles in cell polarity, morphogenesis, migration, and cytokinesis (Pollard and Wu 2010, Cheffings, Burroughs et al. 2016, Misu, Takebayashi et al. 2017, Rottner, Faix et al. 2017, Skruber, Read et al. 2018). A balance of factors that control filament nucleation and stability and those that promote its disassembly exquisitely regulates the functions of the actin cytoskeleton (Lee and Dominguez 2010). Over the past three decades, the fission yeast *Schizosaccharomyces pombe* has emerged as an attractive organism for the study of the actin cytoskeleton and its role in cell polarity and division (Pollard and Wu 2010, Cheffings, Burroughs et al. 2016, Chiou, Balasubramanian et al. 2017). This is particularly due to the fact that many of the actin cytoskeletal proteins can be characterized through easily identifiable lethal mutant phenotypes (Wertman, Drubin et al. 1992, Holtzman, Wertman et al. 1994, Balasubramanian, McCollum et al. 1998).

In fission yeast, the Arp2/3 complex nucleates actin patches (Pelham and Chang 2001, Sirotkin, Berro et al. 2010), whereas linear actin cables and cytokinetic actomyosin rings are assembled by formins For3 (actin cables) (Feierbach and Chang 2001) and Cdc12p (cytokinetic actomyosin rings) (Chang, Drubin et al. 1997) and the actin-binding protein Cdc3-profilin (Balasubramanian, Hirani et al. 1994). The coiled-coil actin binding protein Cdc8 tropomyosin ensures stability of actin cables and actomyosin rings (Liu and Bretscher 1989, Balasubramanian, Helfman et al. 1992, Gunning, Hardeman et al. 2015). The fission yeast actin cytoskeleton undergoes dramatic disassembly and turnover (Kovar, Sirotkin et al. 2011). Treatment of cells with the actin polymerization inhibitor latrunculin A causes complete loss of actin cables, actomyosin rings, and actin patches (in that order) (Pelham and Chang 2001, Pelham and Chang 2002). While the actin severing protein Adf1 (cofilin) plays a key role in actin cytoskeletal disassembly (Nakano and Mabuchi 2006, Pavlov, Muhlrad et al. 2007), the fact that the actin cables and rings still turnover, albeit slowly, in *adf1-1* mutants (Nakano and Mabuchi 2006) suggests that other mechanisms should also exist to disassemble the actin cytoskeleton.

Myo2-S1 (Myo2 G515D) was isolated in a genetic screen for suppressors of the high temperature lethality of *cdc3-124* that led to characterization of the Arp2/3 complex protein Sop2 (Balasubramanian, Feoktistova et al. 1996, Wong, Naqvi et al. 2000), although Myo2-S1 mutant has not been previously characterized. Here we characterize the cellular and molecular basis of the suppression of *cdc3*-124 and other conditions that cause partial loss of the actin cytoskeleton. These studies point to an in vivo role for myosin II in actin filament disassembly.

## Results and Discussion

### *myo2-S1* suppresses the ts lethality of *cdc3-124*

Previous work has shown that *cdc3*-124 is defective in actomyosin ring assembly and colony formation above 32°C (Balasubramanian, Feoktistova et al. 1996). To investigate and recreate the suppression of *cdc3*-124 by *myo2*-S1, we freshly generated *cdc3*-124 *myo2*-S1 from a cross between a *myo2*-S1 parent (obtained after three rounds of back crossing with wild-type cells) and *cdc3*-124. The newly created *cdc3*-124 *myo2*-S1 strain was compared to wild-type and to *cdc3*-124 and *myo2*-S1 single mutants. As expected, wild-type cells formed colonies at all temperatures tested, whereas *cdc3*-124 failed to form colonies at and above 32°C (Figure 1A). *myo2*-S1 was able to form colonies at all temperatures tested, although these cells displayed cytokinesis defects at all temperatures tested (described in later sections) (Figure 1A). Importantly, the recreated *cdc3*-124 *myo2*-S1 was able to form colonies at 34°C and did so poorly, even at the higher restrictive temperature of 36°C (Figure 1A). The genetic screen (Balasubramanian, Feoktistova et al. 1996) also identified a different allele of *myo2*, *myo2*-S2 (Myo2 E679K)(Wong, Naqvi et al. 2000), which rescued *cdc3*-124 (Figure 1A). We recreated the *cdc3*-124 *myo2*-S2, using a back-crossed *myo2*-S2 parent, and found that it rescued *cdc3*-124 at 34°C, but barely rescued it at 36°C (Figure 1A). The well-characterized *myo2*-E1 allele (Balasubramanian, McCollum et al. 1998) did not rescue *cdc3*-124 either at 34°C or 36°C. Since *myo2*-S1 better suppressed the *cdc3*-124 defect, we characterized this mutant further in the rest of this study.

**Figure 1:**
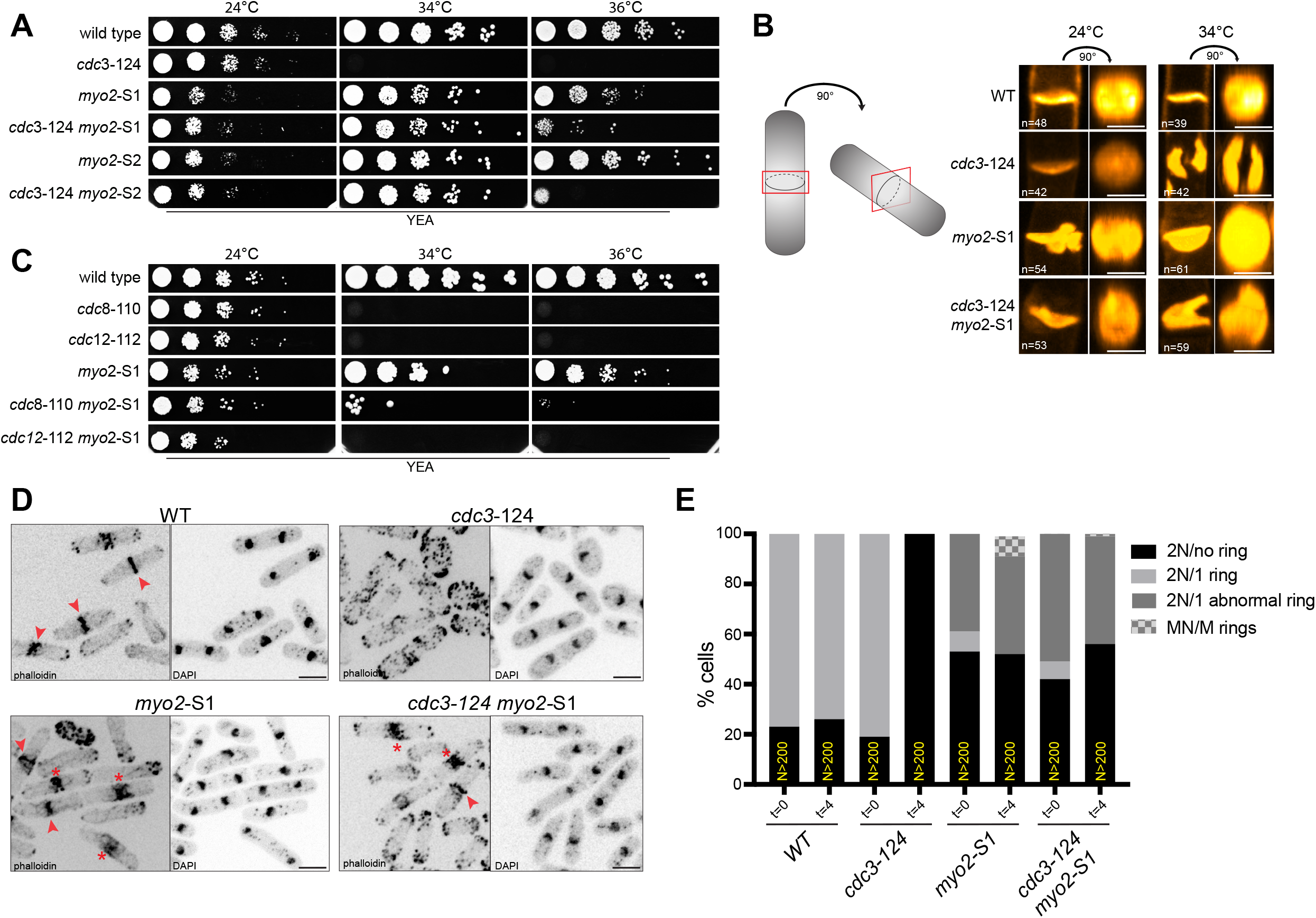
*myo*2-S1 suppresses the lethality of *cdc3-124* at the non-permissive temperature. A) 10-fold serial dilutions of wild-type, *cdc3*-124, *myo2*-S1, *cdc3*-124 *myo2*-S1, *myo2-S2* and *cdc3*-124 *myo2-S2* were spotted onto YEA agar plates and grown for 3 days at 24°C, 34°C and 36°C. B) Calcofluor white staining was used to visualize the septum of wild-type, *cdc3*-124, *myo2*-S1 and *cdc3*-124 *myo2-S* cells, fixed at 24°C or after 4 hours shift at 34°C. In the first column it is shown the acquired front view image of the septum, while in the second column it’s displayed the face-on view of the septum, which was generated with Fiji software (as illustrated by the cartoon). Scale bar represent 3 μm. C) 10-fold serial dilutions of wild-type, *cdc8*-110, *cdc12-112, myo2-S1, cdc8-110 myo2-S1* and *cdc12-112 myo2-S1* were spotted onto YEA agar plates and grown for 3 days at 24°C, 34°C and 36°C. D) Cells were grown at 24°C and shifted for 4 hours at 34°C before PFA fixation. CF633-phalloidin and DAPI were used to visualize actin structures and the nucleus respectively, of wild-type, *cdc3*-124, *myo2*-S1and *cdc3*-124 *myo2*-S1. Arrows indicated normal actomyosin rings while asterisks indicated abnormal rings. Scale bar represent 5 μm. E) Quantification of CF633-phalloidin and DAPI staining in (D) is shown. Cells with 2 nuclei were classified depending on the presence of either no ring (2N/no ring), 1 normal ring (2N/1 ring), 1 abnormal ring (2N/abnormal ring) or multiple nuclei and rings (MN/M rings).

The strains described above were grown in liquid medium and stained with calcofluor white to visualize the division septa. Whereas normal (or nearly normal) division septa, that appeared as a disc in tilted images were detected at 24°C in all strains, complete septa that appeared as a disc in tilted images were found in wild-type, *myo2*-S1, and *cdc3*-124 *myo2*-S1, but not in *cdc3*-124, at 34°C (Figure 1B). Instead, patches of septum material that did not create a barrier between the two daughters were observed in *cdc3*-124 at 34°C (Figure 1B). This experiment established that compromise of *myo2*, through the *myo2*-S1 mutation, led to full rescue of the septation defect of *cdc3*-124.

Cdc3-profilin (Balasubramanian, Hirani et al. 1994), in conjunction with Cdc12-formin (Chang, Drubin et al. 1997) plays a key role linear actin filament nucleation during cytokinesis (Kovar, Kuhn et al. 2003), whereas Cdc8-tropomyosin (Balasubramanian, Helfman et al. 1992) plays a role in actin cable and ring stabilization (Gunning, Hardeman et al. 2015, Khaitlina 2015). We therefore tested if the observed suppression was specific to Cdc3-profilin defects or if *myo2*-S1 also suppressed defects in *cdc12* and *cdc8.* To test this, we generated double mutants *cdc8*-110 *myo2*-S1 and *cdc12-112 myo2-S1.* We found that *cdc8*-110 was suppressed at 34°C and tiny “pin-prick” colonies were observed even at 36°C (Figure 1C). *cdc12-112* was not suppressed at either 34°C or at 36°C (Figure 1C).

Next, we stained wild-type, *cdc3*-124, *myo2*-S1 and *cdc3*-124 *myo2*-S1 with CF633-phalloidin and DAPI to observe the actin cytoskeleton and nuclei (Figure 1D and E). This analysis showed that whereas actin rings were not observed in *cdc3*-124, actin rings were observed in mitotic wild-type, *myo2*-S1, and *cdc3*-124 *myo2*-S1 mutants (Figure 1D). The rings in *myo2*-S1 were more diffuse compared to those in wild-type consistent with a role for Myo2p in actomyosin ring assembly. These experiments established that compromise of Myo2p function led to suppression of the defective actomyosin ring assembly, septum assembly, and colony formation in *cdc3*-124 (Figure 1E). The partial suppression of *cdc8*-110 by *myo2*-S1 (Figure 1C), suggested that *myo2*-S1 suppression is not specific to *cdc3*-124 and that it also suppress defects in other actin assembly / stability factors.

To investigate the mechanism of suppression of *cdc3*-124 by *myo2*-S1, as well as to characterize the phenotypic consequences of *myo2*-S1, if any, we generated wild-type, *cdc3*-124, *myo2*-S1, and *cdc3*-124 *myo2*-S1, expressing Rlc1-3GFP (actomyosin ring marker) and mCherry-Atb2 (mitotic spindle marker). At the permissive temperature of 24°C, actomyosin ring assembly and contraction were indistinguishable in wild-type and *cdc3*-124 cells, whereas both processes were slightly slower in *myo2*-S1 and *cdc3*-124 *myo2*-S1 cells (Figure S1A and B). Ring assembly in wild-type at 34°C took 17 ± 6.8 minutes and ring contraction took 18.7 ± 2.1 minutes (Figure 2B and C). As expected, actomyosin rings did not assemble in *cdc3*-124 cells although Rlc1-3GFP accumulated first in nodes and then in multiple spots and bar-like structures (Figure 2A). The *myo2*-S1 mutant showed a range of phenotypes pertaining to cytokinesis. First, in ~ 50% of the cells, ring assembly was delayed and took ~ 1.8 times longer (30.2 ± 4 minutes) than in wild-type cells and fully compacted rings were observed only in cells with elongated mitotic spindles (Figure 2A class I and class III and Figure 2B). Second, the remaining ~ 50% of the cells made abnormal actomyosin structures that did not compact into a ring structure (Figure 2A class IV). Ring contraction was uniformly slower in cells that appeared to have normal rings (38.3 ± 6.9 minutes) (Figure 2C). Furthermore, ring contraction was asymmetric in nearly two-thirds of cells that appeared to have a normal looking ring (Figure 2A class III). The *cdc3*-124 *myo2*-S1 double mutant largely resembled *myo2*-S1 single mutant and rings of normal appearance were detected in ~ 42% of the cells and these rings took 26.2 ± 1.9 minutes to assemble, while the rest of the cells assembled Rlc-3GFP bundles that did not compact into a ring (Figure 2A and B). Ring contraction in *cdc3*-124 *myo2*-S1 cells was almost always asymmetric and the process took almost three times the time taken in wild-type cells (56.3 ± 26.8 minutes) (Figure 2A and C). These experiments established that *myo2*-S1 was compromised for both known physiological roles of Myo2p, i.e. actomyosin ring assembly and contraction, and that the compromise of one of its known activities contributed to the genetic suppression of cytokinesis and colony formation defects of *cdc3*-124.

**Figure 2:**
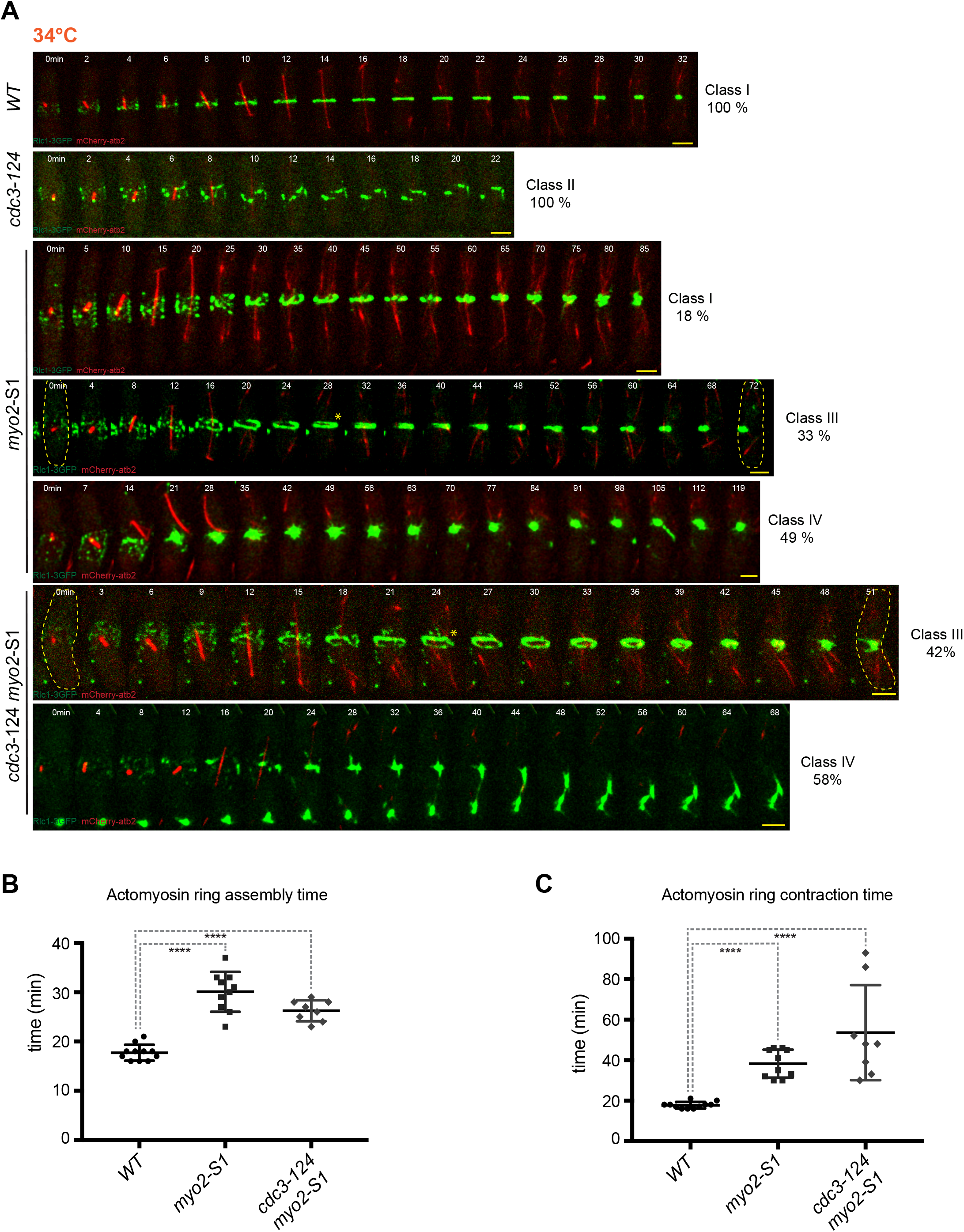
Actomyosin ring assembly and contraction is partially restored in *myo2*-S1 and *cdc3*-124 *myo2*-S1 at 34°C. A) Time-lapse series of wild-type, *cdc3*-124, *myo2*-S1and *cdc3*-124 *myo2*-S1 cells expressing Rlc1-3GFP as a contractile ring marker and mCherry-atb2 as a cell-cycle stage marker. Cells were grown at 24°C and shifted 3 hours at 34°C before being imaged at 34°C. More than 20 cells were imaged and quantified for each strain. On the site of each montage it is indicated the percentage of the different cytokinetic behaviours categorized into class I (normal actomyosin ring – AMR), class II (failed AMR assembly), class III (normal AMR assembly and asymmetrical AMR contraction) and class IV (abnormal AMR structures). Images shown are maximum-intensity projections of Z-stacks. Time indicated in minutes. Scale bars represent 3 μm. B) Quantification of the time necessary for actomyosin ring assembly of class I and class III cells imaged in (A). Statistical significance was calculated by Student’s test (****p<0.0001). Error bars represent SD. C) Quantification of the time necessary for actomyosin ring contraction of class I and class III cells imaged in (A). Statistical significance was calculated by Student’s test (****p<0.0001). Error bars represent SD.

### ATP-dependent contraction is dramatically slowed in actomyosin rings isolated from myo2– S1 cells

The mutated myosin allele, Myo2-S1, possesses a point mutation resulting in the replacement of Gly515 with an aspartate in the L50 subdomain of Myo2p motor domain. G515D is located at the beginning of HR helix just at the end of activation loop (Varkuti, Yang et al. 2012). This glycine might play a role in the conformational flexibility of the activation loop, thereby contributing to the loop’s orientation that facilitates actin binding. In order to understand the effect of the mutation, we analyzed the position equivalent to G515 in various myosin structures. In the cryoEM structure of myosin bound to actin filaments (5JLH), G515 was observed to be at the beginning of the activation loop that interacted with N-terminal end of an actin monomer (Figure 3A). Modeling of the G515D mutation on myosin motor domain structure (1VOM) showed a salt bridge formation between the mutated Asp515 and Arg509 residues (data not shown). Arg520 in *C. elegans* muscle myosin, corresponding to Lys510 in *S. pombe* Myo2p, has been shown to be crucial for binding with the N-terminal region of actin (Varkuti, Yang et al. 2012). Furthermore, Arg520 mutation affected the ATPase activity of myosin motor domain indicating that Arg520 interaction with actin is essential for motor domain function (Varkuti, Yang et al. 2012). In a sequence alignment of 237 non-redundant myosin sequences, glycine is highly conserved and a positively charged residue (at least one arginine or lysine) is a conserved feature of the activation loop (Figure 3B). Unavailability or misorientation of the positively charged side chain (Lys or Arg) to interact with actin in the G515D mutant (Myo2-S1) might weaken its interaction with actin, leading to abnormalities in ring assembly. The mutation might also affect the motor activity in the defective rings due to the absence of ATPase stimulation upon actin interaction, thus affecting the actomyosin ring contraction rates.

**Figure 3:**
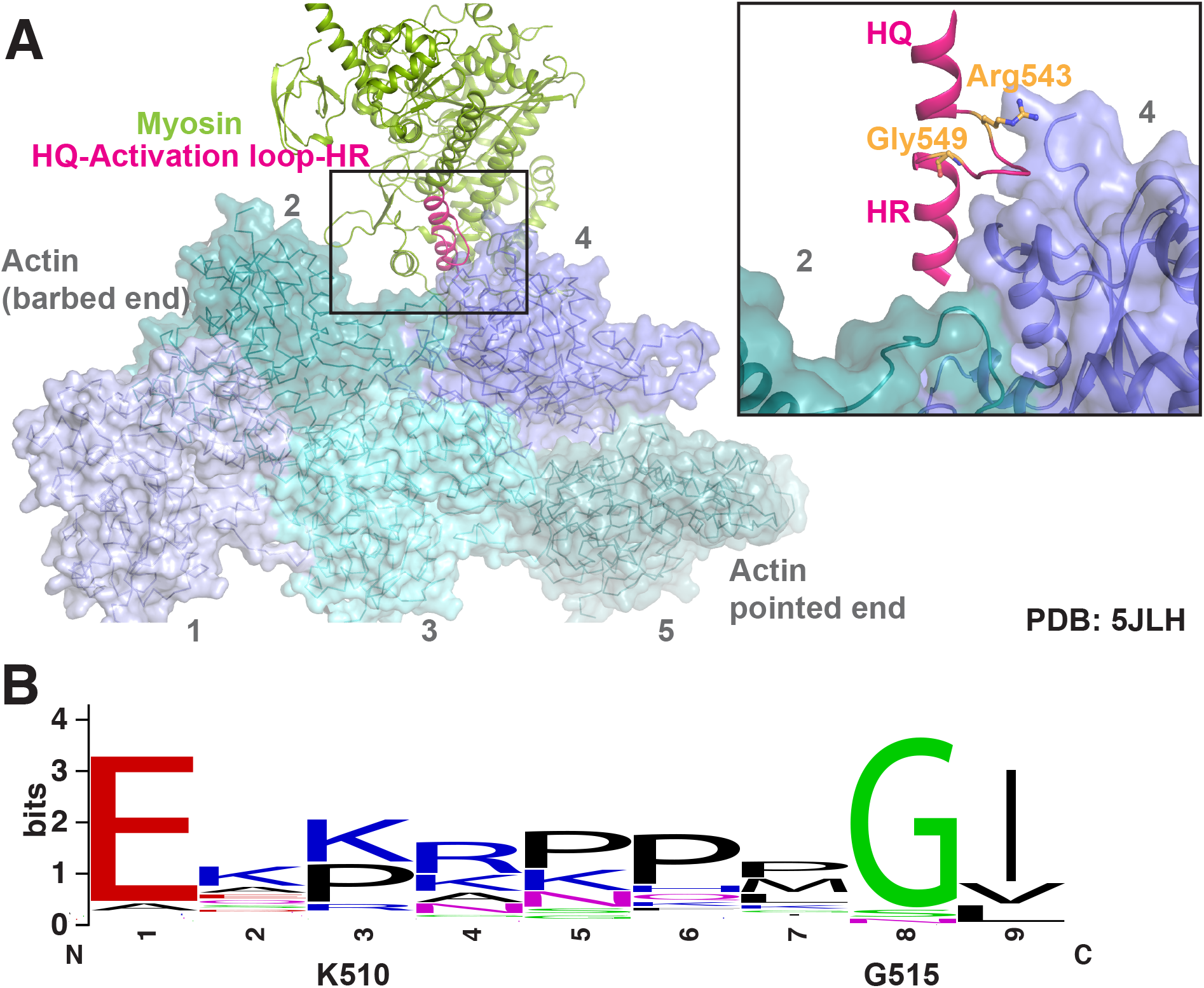
Structural basis of *S. pombe* Myo2-S1 (G515D) mutation. A) Gly515 in *S. pombe* Myo2 (equivalent to Gly549 of human myosin 14 in the actomyosin complex structure of PDB ID: 5JLH) is positioned at the boundary of the activation loop at the myosin-actin interface. Inset shows the zoomed view of the interaction between the activation loop and the N-terminus of actin protomer. Gly549 and Arg543, corresponding to Gly515 and Lys510 of *S. pombe* Myo2, are shown in stick representation. The figure was generated using PyMol. B) Conservation information of Gly515 and Lys510 depicted on a sequence logo representation. The figure was generated using WebLogo (Crooks, Hon et al. 2004) using a sequence alignment of 237 non-redundant myosin sequences aligned using ClustalO (Sievers, Wilm et al. 2011, Sievers and Higgins 2018). The sequence corresponding to the activation loop was selected using JalView (Waterhouse, Procter et al. 2009).

Given these considerations based on structural analysis of *myo2*-S1, we wanted to test the biochemical properties of Myo2-S1p, especially as it pertains with actin binding or motor activity. We were unable to purify this protein in sufficient quantities for biochemical studies (data not shown). As an alternative, we therefore used the permeabilized spheroplast assay, in which actomyosin rings contract in an ATP and myosin II dependent manner (Mishra, Kashiwazaki et al. 2013, Huang, Mishra et al. 2016). We prepared cell ghosts from wild-type, *myo2-S1* and *cdc3-124 myo2-S1* cells expressing Rlc1-3GFP that were grown at 24°C. We carried out two sets of experiments, one at 24°C (Figure S2A) and the other at 34°C (Figure 4A and S2B). Actomyosin rings in wild-type cell ghosts were stably maintained and contracted rapidly upon ATP addition at 24°C (Figure S2A and 4C). However, actomyosin rings became extremely unstable upon introduction of the *myo2*-S1 mutation, suggesting that Myo2-S1p may interact weakly with actin causing the ring to be unstable in the context of a cell ghost (Figure S2A). We therefore stabilized actomyosin rings by incubation of cell ghosts with the actin-stabilizing compound Jasplakinolide (Jas) (Holzinger 2009). We next treated Jas-stabilized actomyosin rings in cell ghosts with ATP. As observed in the absence of Jas, rings in wild-type ghosts contracted rapidly at 34°C (Figure 4A and B). However, even though actomyosin rings were stably maintained in *myo2*-S1 and *cdc3*-124 *myo2*-S1, they either failed to contract or were extremely slow to contract (Figure 4A and B). A similar trend was observed when Jas-stabilized wild-type, *myo2*-S1, and *cdc3*-124 *myo2*-S1 ghosts were incubated with ATP even at the lower temperature of 24°C (Figure 4C). This is similar to our previous observations with cell ghosts of the well characterized temperature-sensitive *myo2-* E1 allele, rings from which fail to contract upon ATP addition, even at the permissive temperature of 24°C (Mishra, Kashiwazaki et al. 2013, Palani, Chew et al. 2017). In addition, we found that Jas-stabilized actomyosin rings in cell ghosts prepared from *cdc3*-124 *myo2-S2* (which suppresses *cdc3*-124, but less well compared to *myo2*-S1) also underwent very slow ATP-dependent contraction at 34°C (Figure S2B and C) and at 24°C (Figure S2D). These observations established that loss of myosin II motor activity strongly correlated with the suppression of actomyosin ring assembly / stability defect in *cdc3*-124. The fact that two different mutants, which show defects in ATP-dependent ring contraction *in vitro,* suppress *cdc3*-124 suggests that reduced actin binding and / or motor activity rather than some other defect in Myo2p contributes to the rescue of *cdc3*-124.

**Figure 4:**
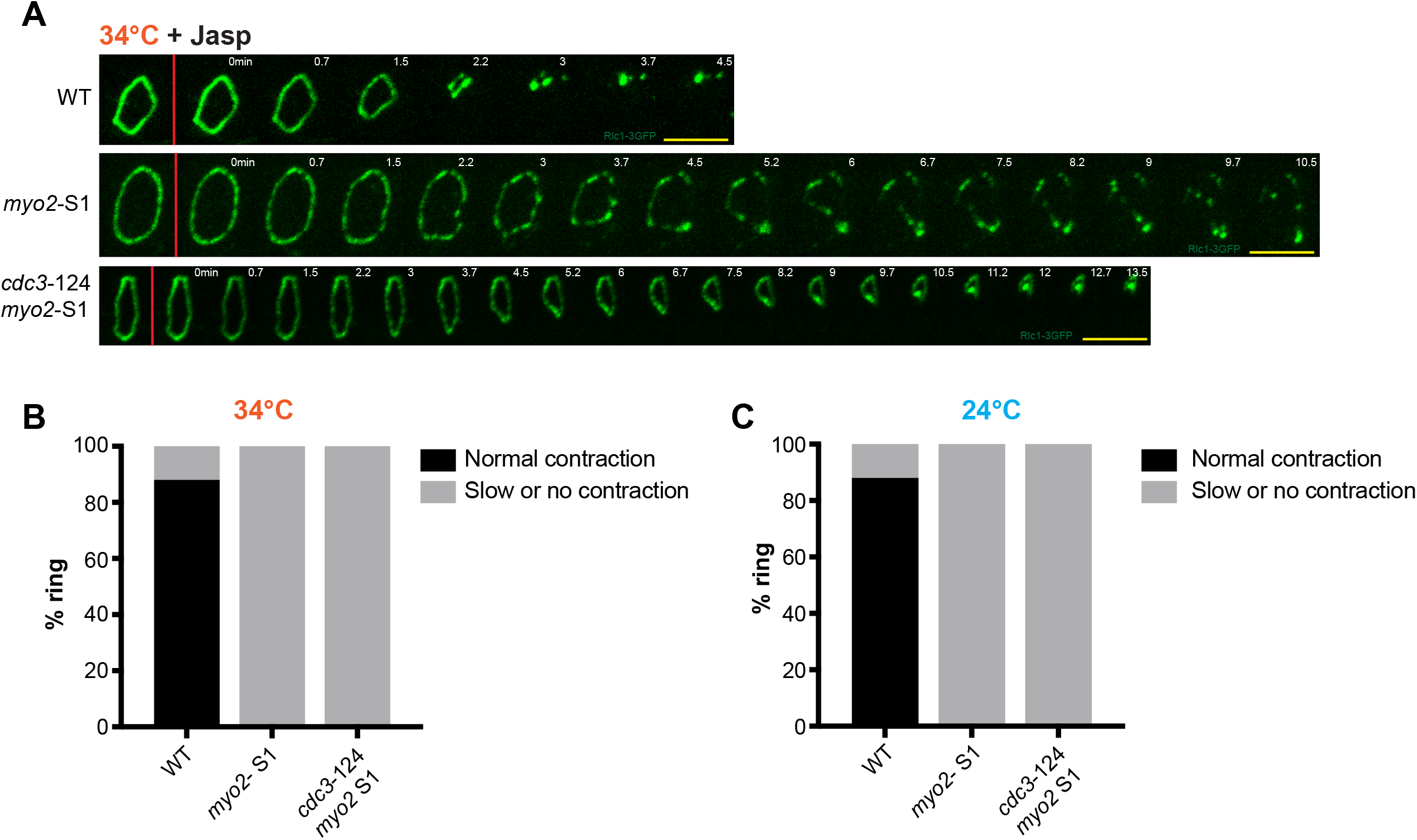
Actomyosin rings isolated from *myo2-S1* and *cdc3-124 myo2-S1* cells do not undergo ATP-dependent contraction. A) *In vitro* isolated actomyosin rings were prepared from wild-type (n=17), *myo2*-S1 (n=14), and *cdc3*-124 *myo2*-S1 (n=20) cells grown at 24°C in the presence of 20 μM jasplakinolide (jasp). Isolated rings were shifted 15 minutes at 34°C before proceeding with imaging. Ring contraction experiments were performed at 34°C and contraction was activated by addition of 0.5 mM ATP (indicated by the red bar). Images shown are maximum intensity projections of Z-stacks. Time indicated in minutes. Scale bar represent 5 μm. B) Percentage of rings that either contracted normally or presented slow/no contraction of strains imaged at 34°C illustrated in (A). C) Percentage of rings that either contracted normally or presented slow/no contraction of strains imaged at 24°C illustrated in (A).

### *myo2-S1* suppresses direct pharmacological perturbation of the actin cytoskeleton with Latrunculin A

Previous work has shown that myosin II can break actin filaments *in vitro* both by stretching or buckling (Murrell and Gardel 2012, Vogel, Petrasek et al. 2013) due to its motor activity. We have shown that myosin II mutants defective in ATP dependent contraction (and by inference in motor activity) suppress defects caused by partial loss of proteins contributing actin cable and ring assembly / stability. We reasoned that shorter or unstable actin filaments may be protected for longer periods in the *cdc3*-124 *myo2*-S1, *cdc8*-110 *myo2*-S1, and *cdc3*-124 *myo2-S2* mutants, which in turn may contribute to the observed suppression of the cytokinesis and colony formation defects. We therefore tested if direct pharmacological perturbation of actin cables and actomyosin rings (which are composed of formin generated linear actin filaments) by treatment with low doses of Latrunculin A (Ayscough, Stryker et al. 1997) can be reversed by compromise of Myo2p. Previously, we have shown that treatment of wild-type cells with low doses of Latrunculin A (~ 0.25 μM) leads to cytokinesis defects (Mishra, Karagiannis et al. 2004). We plated serial dilutions of wild-type, *cdc3*-124, *myo2*-S1, and *cdc3*-124 *myo2*-S1 with a series of doses of Latrunculin A and incubated these at 30°C and 34°C (Figure 5A). We found that wild-type cells were capable of robust colony formation up to 0.25 μM Latrunculin A, but did not form colonies at 0.375 μM Latrunculin A (Figure 5A). Consistent with a role for Cdc3p profilin in actin cable and actomyosin ring assembly, *cdc3*-124 mutants were hypersensitive to 0.25 μM Latrunculin A. Interestingly, *myo2*-S1 mutants were able to form robust colonies at all temperatures tested up to 0.375 μM Latrunculin A (Figure 5A), establishing that compromise of Myo2p function reverses the deleterious effects caused by actin perturbation. The *cdc3*-124 *myo2*-S1 was also capable of colony formation at 30°C at 0.375 μM Latrunculin A, although at 34°C, this strain only grew on plates containing 0.125 μM Latrunculin A (Figure 5A). The inability of the *cdc3*-124 to form colonies at and above 0.25 μM Latrunculin A potentially reflects an enhanced actin cytoskeletal defect due to the additive effect of loss of Cdc3p and Latrunculin A. CF633-Phalloidin staining of liquid cultures of the four strains described above showed that actin cables, rings, and patches were present at 0 time point in all four strains (Figure 5B and C). After a 3 hours incubation in 0.375 μM Latrunculin A, actin patches were observed in all 4 strains. However, actin cables / rings were not detected in wild-type or *cdc3*-124 but were clearly detected in *myo2*-S1 and *cdc3*-124 *myo2*-S1 cells (Figure 5B and C). Collectively, these experiments established that partial genetic or pharmacological perturbation of actin cables and rings can be suppressed by motor-activity defective Myo2p.

**Figure 5:**
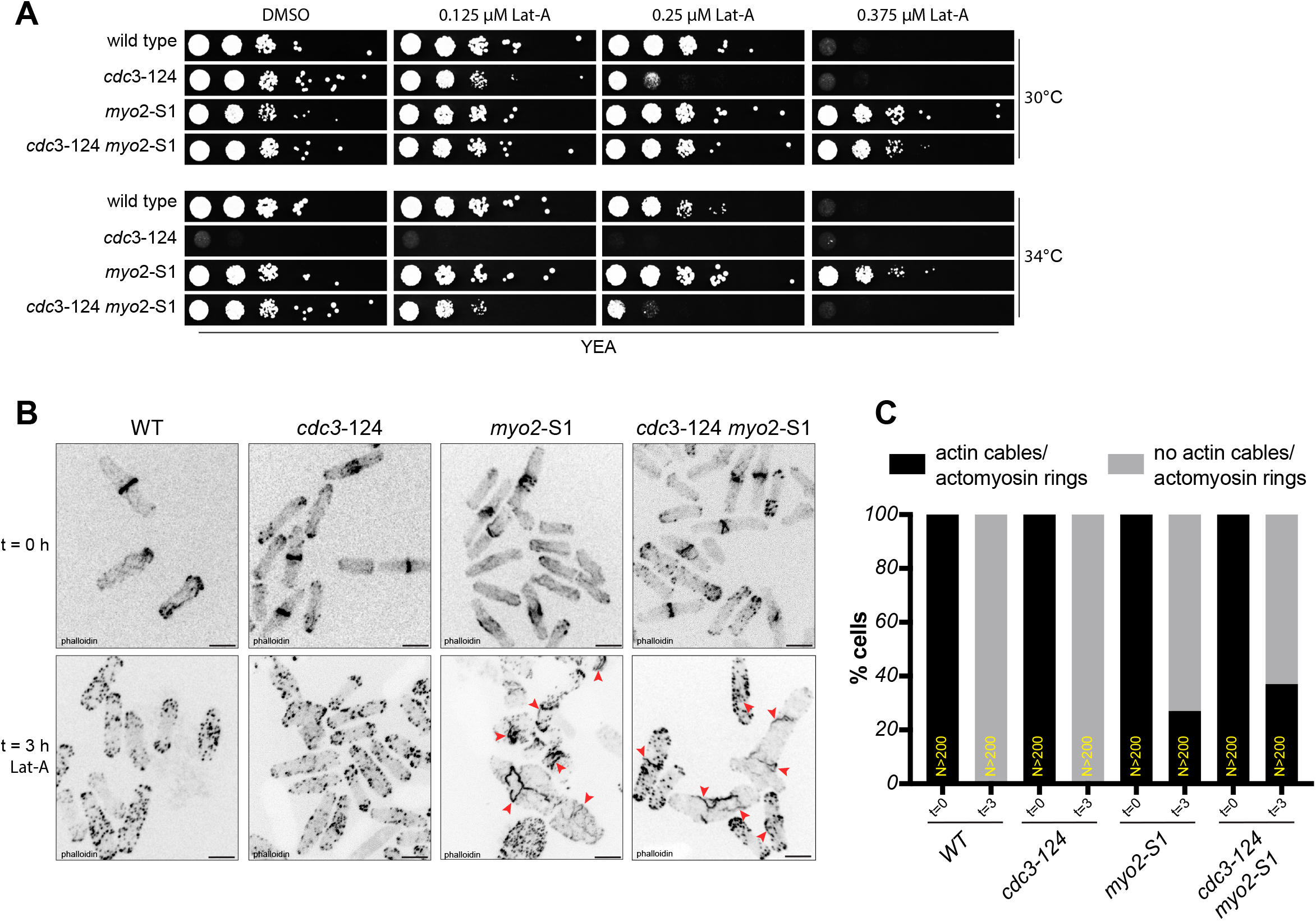
myo2-S1 suppresses perturbations of actin induced by Latrunculin A. A) 10-fold serial dilutions of wild-type, *cdc3*-124, *myo2*-S1and *cdc3*-124 *myo2*-S1 cells were spotted onto YEA agar plates containing different concentrations of latrunculin-A (Lat-A) and grown for 3 days at 24°C, 34°C and 36°C. B) Cells were grown at 24°C and shifted for 3 hours at 34°C in the presence of 0.375 μM of Lat-A before PFA fixation. CF633-phalloidin was used to visualize actin structures of wildtype, *cdc3*-124, *myo2*-S1 and *cdc3*-124 *myo2*-S1 cells. Arrows indicated the presence of actin cables. Scale bar represent 5 μm. C) Quantification of the presence of actin cables or actomyosin ring structure detected with CF633-phalloidin staining in (D). More than 200 cells were imaged for wild-type, *cdc3*-124, *myo2*-S1and *cdc3*-124 *myo2*-S1.

In summary, in this study we provide evidence for a role for myosin II in actin filament disassembly through the use of genetic, structural, and *in vitro* analyses. Previous work has shown that single actin filaments and networks of actin filaments are broken and disassembled by myosin II through buckling (Murrell and Gardel 2012, Vogel, Petrasek et al. 2013). Previous work in *S. japonicus* has shown that isolated cytokinetic rings break into a series of clusters in a myosin II dependent manner, which is reversed and normal contraction upon ATP addition ensues, when the isolated cytokinetic rings are pre-incubated with Jasplakinolide (Chew, Huang et al. 2017). Our work may provide *in vivo* evidence for a role for myosin II in actin filament disassembly and turnover, also consistent with the work in cytokinetic mammalian cells (Murthy and Wadsworth 2005). It is possible that formin-generated actin filaments are short and unstable in profilin and tropomyosin mutants and upon exposure of cells with low doses of Latrunculin A. In this scenario, compromise of myosin II, which normally destabilizes actin filaments, rescues the instability of actin filaments and, in turn, the cytokinesis defects in *cdc3*-124, *cdc8*-110 and Latrunculin A treated cells.

## Materials and methods

### Yeast genetics and culture methods

Cells were grown and cultured at 24 °C in yeast extract medium (YES) as described previously (Moreno, Klar et al. 1991). The presence of designated mutations on each used strain has been verified by PCR and DNA sequencing.

### PFA fixation and fluorescence microscopy

*S. pombe* cells, growing at 24°C in YES medium, were either fixed in mid-log phases or shifted to 34°C for 3-4 hours before fixation. Cells were fixed in a 4% paraformaldehyde solution for 12 minutes at room temperature and, after two washes with 1X PBS, permeabilized with 1% Triton X100 for 10 minutes. Cells were washed twice with 1X PBS and stained with either DAPI (4’,6-diamidino-2-phenylindole, from Life Technologies), for the visualization of the nucleus, or CF633-Phalloidin (CF633, from Biotium Inc) to visualize actin structures. To visualize the septa, fixed *S. pombe* cells were directly incubated with Calcofluor white (CW, from Sigma-Aldrich). Still images were acquired using a spinning disk confocal microscope (specifications described below in livecell imaging section) and processed using imaging software Fiji.

### Live-cell imaging

Mid-log phase cells were grown at 24°C in YES medium and, when necessary, shifted at 34°C for 3-4 hours before imaging acquisition of live cells in a temperature-controlled incubation chamber. Time-lapse images were acquired for 3-4 hours using a spinning disk confocal microscope (Andor Revolution XD imaging system, equipped with a 100x oil immersion 1.45NA Nikon Plan Apo lambda, and a confocal unit Yokogawa CSU-X1, EMCCD detector (Andor iXON) and Andor iQ acquisition software). Cells were imaged in a CellASIC microfluidic yeast plates (Y04C and D size), where 15 Z-stacks of 0.5 μm thickness images were acquired at 1-minute intervals for Rlc1-3GFP (myosin regulatory light chain, used as actomyosin ring marker) and Atb2-mCherry (alpha tubulin 2, used as cell cycle marker). PRISM 6.0 software (GraphPad) was used for quantification and the statistical significance was determined using Student’s t-test (****P<0.0001).

### Isolation of actomyosin rings and ATP-dependent contraction

Actomyosin rings were isolated as described previously (Mishra, Kashiwazaki et al. 2013, Huang, Mishra et al. 2016). Time-lapse imaging of Rlc1-3GFP were acquired as described before in a CellASIC microfluidic yeast plates (Y04D), where 21 Z-stacks of 0.5 μm thickness images were acquired while treating the cells with 0.5 mM ATP, either at 24°C or after shifting 10 minutes the isolated ring at 34°C. Fiji imaging software was used to process the acquired images, obtaining the maximum intensity projection of the Z-stacks.

### Drug treatments in cells and isolated actomyosin rings

YES agar plates in Figure 5A were prepared by the addition of different concentrations of latrunculin A (Lat-A; Enzo Life Sciences), where cells were successively being spotted at 10-fold serial dilutions. For the treatment in Figure 5B, cells were grown in liquid YES medium at 24°C and successively shifted at 34°C, in the presence of 0.375 μM Lat-A in the medium, for 3 hours before fixation.

Isolated actomyosin rings were treated, when indicated, with jasplakinolide (jasp; Enzo Life Sciences) by the addition of the drug to the isolated rings at the final concentration of 20 μM.

### Table of strains used in this study

|  |  |
| --- | --- |
| MBY17 | *cdc3-124 leu1-32 ura4-D18 h+* |
| MBY54 | *myo2-S2 ade6-21 ura4-D18 h-* |
| MBY53 | *myo2-S1 ade6-21 ura4-D18 leu1-32 h-* |
| MBY109 | *cdc8-110 ade6-210 ura4-D18 leu1-32 h-* |
| MBY144 | *cdc12-112 ura4-D18 leu1-32 h-* |
| KGY247 | *ade6-210 ura4-D18 leu1-32 h+* |
| MBY12728 | *myo2-S1 cdc3-124 ade6-21 ura4-D18 leu1-32* |
| MBY12730 | *myo2-S1 rlc1-3GFP::kanMx6 mCherry-atb2::hph ade6-21 ura4-D18 leu1-32 h+* |
| MBY12732 | *myo2-S1 cdc3-124 rlc1-3GFP::kanMx6 mCherry-atb2::hph ade6-21 ura4-D18 leu1-32 h-* |
| MBY12734 | *myo2-S2 cdc3-124 ade6-21 ura4-D18 leu1-32* |
| MBY12736 | *myo2-S2 rlc1-3GFP::kanMx6 mCherry-atb2::hph ade6-21 ura4-D18 leu1-32* |
| MBY12738 | *myo2-S2 cdc3-124 rlc1-3GFP::kanMx6 mCherry-atb2::hph ade6-21 ura4-D18 leu1-32 h+* |
| MBY12928 | *myo2-S1 cdc8-110 ade6-21 ura4-D18 leu1-32* |
| MBY12930 | *myo2-S1 cdc12-112 ade6-21 ura4-D18 leu1-32* |
| MBY12931 | *myo2-S2 cdc8-110 ade6-21 ura4-D18 leu1-32* |
| MBY12933 | *myo2-S2 cdc12-112 ade6-21 ura4-D18 leu1-32* |

## Acknowledgements

This work was funded by research grants from Wellcome Trust (WT101885MA) and the European Research Council (GA 671083 – ACTOMYOSIN RING) to MKB. PG was supported by research grant from SERB Women Excellence Award and computation facilities supported by the Collaborative Research Grant from CEFIPRA (Indo-French Centre for Promotion of Advanced Research). SJ acknowledges INSPIRE for fellowship. We thank Dr. Bernardo Chapa-y-Lazo for help with image analysis.

## Supplemental Figure Legends

**Supplemental 1**

A) Quantification of the time necessary for actomyosin ring assembly of class I and class III cells imaged at 24°C in wild-type, *cdc3*-124, *myo2*-S1and *cdc3*-124 *myo2*-S1 strains. Statistical significance was calculated by Student’s test. Error bars represent SD.

B) Quantification of the time necessary for actomyosin ring contraction of class I and class III cells imaged at 24°C in wild-type, *cdc3*-124, *myo2*-S1and *cdc3*-124 *myo2*-S1 strains. Statistical significance was calculated by Student’s test. Error bars represent SD.

**Supplemental 2**

A) *In vitro* isolated actomyosin rings were prepared from wild-type (n=17) and *myo2*-S1 (n=15) cells in the absence of jasplakinolide. Ring contraction experiments were performed at 24°C and contraction was activated by addition of 0.5 mM ATP (indicated by the red bar). Images shown are maximum intensity projections of Z-stacks. Time indicated in minutes. Scale bar represent 5 μm.

B) *In vitro* isolated actomyosin rings were prepared from *cdc3*-124 *myo2-S2* cells (n=23) grown at 24°C in the presence of 20 μM jasp. Isolated rings were shifted 15 minutes at 34°C before proceeding with imaging. Ring contraction experiments were performed at 34°C and contraction was activated by addition of 0.5 mM ATP (indicated by the red bar). Images shown are maximum intensity projections of Z-stacks. Time indicated in minutes. Scale bar represent 5 μm.

C) Percentage of rings isolated from wild-type, *myo2-S2* and *cdc3*-124 *myo2-S2* cells that either contracted normally or presented slow/no contraction imaged at 34°C

**D)** Percentage of rings isolated from wild-type, *myo2-S2* and *cdc3*-124 *myo2-S2* cells that either contracted normally or presented slow/no contraction imaged at 24°C.

