## Supplementary figures and images for "Genetic interaction between profilin and myosin II reveals a potential role for myosin II in actin filament disassembly *in vivo*"

### Supplemental Figures

#### Supplement S1

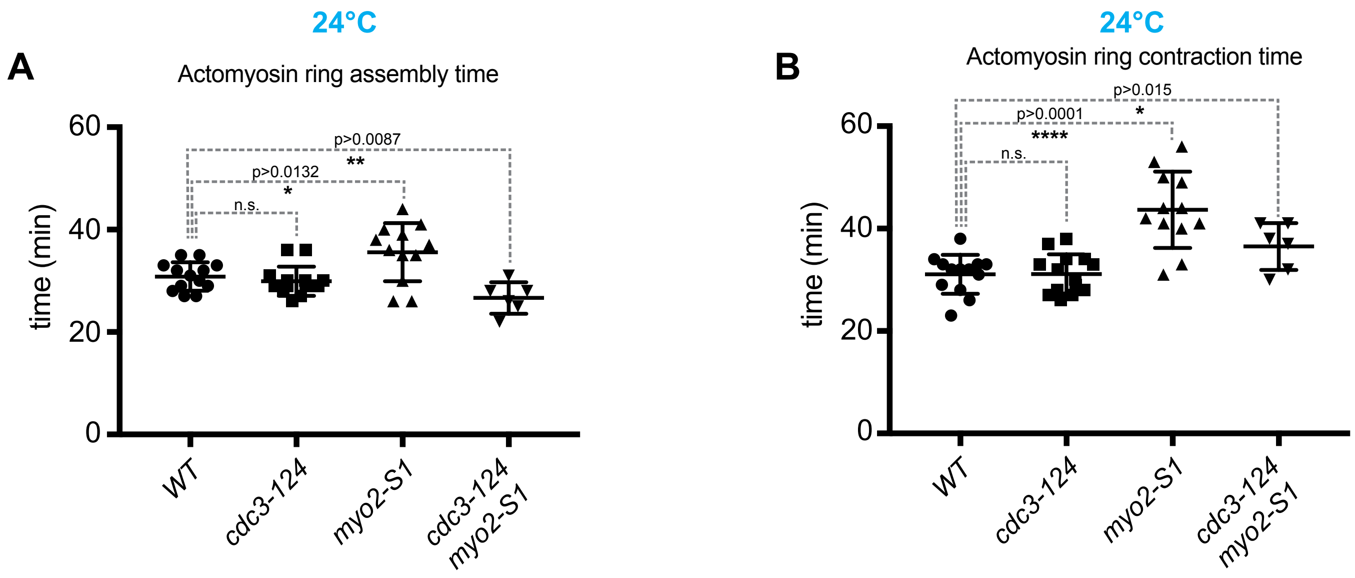

##### Supplement S2

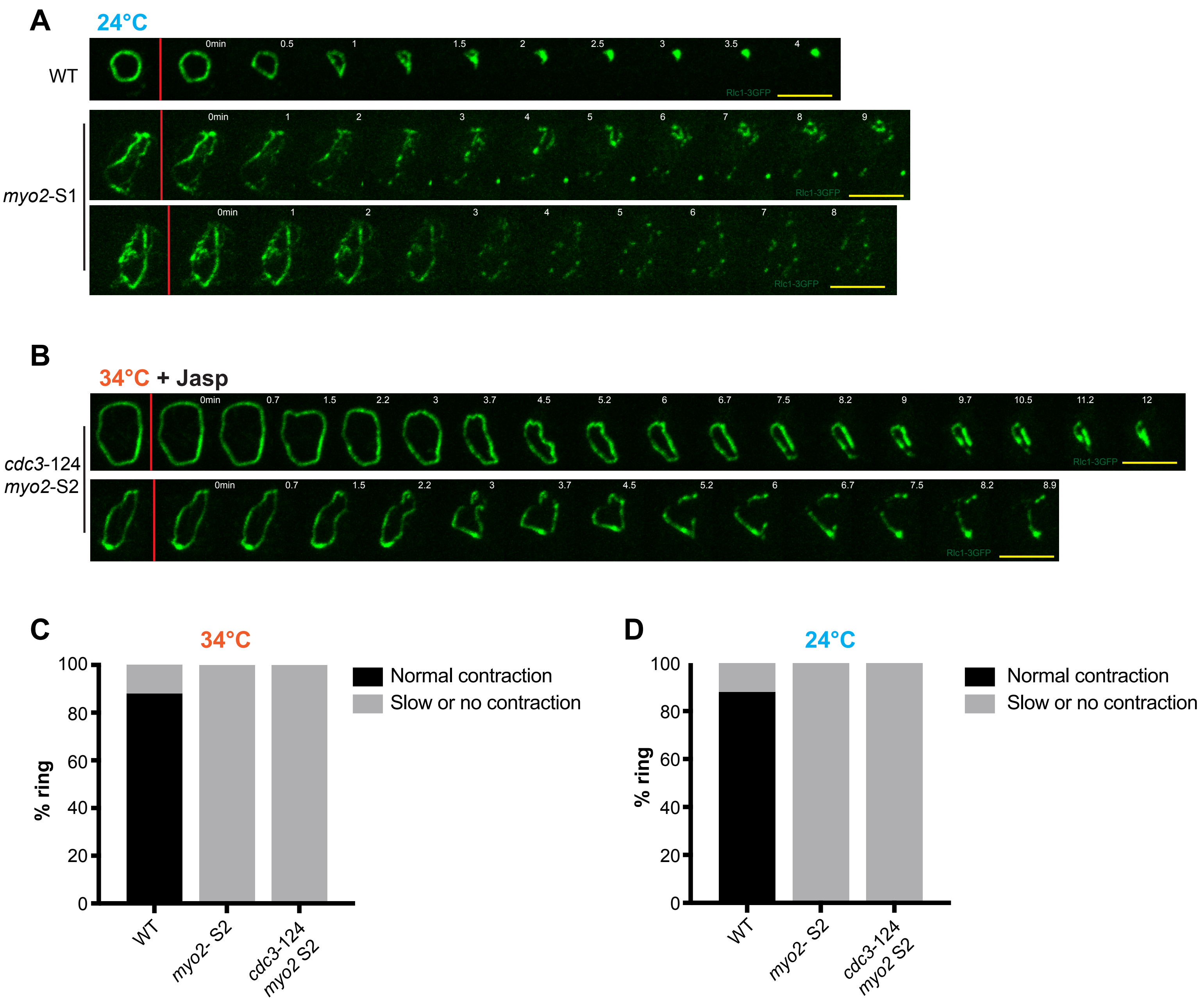
